# Thrombin cleaves and activates the protease-activated receptor 2 dependent on thrombomodulin co-receptor availability

**DOI:** 10.1101/550137

**Authors:** Dorothea M. Heuberger, Alessandro G. Franchini, Jerzy Madon, Reto A. Schuepbach

## Abstract

**Introduction:** Protease-activated receptors (PARs) evolved to react to extracellular proteolytic activity. In mammals, three of the four PARs (PAR1, PAR3, and PAR4) that are expressed respond to the prototypical procoagulant enzyme thrombin, whereas PAR2 was assumed to resist activation by thrombin. To date, involvement of cell surface thrombin-recruiting co-receptors such as thrombomodulin (TM), which potentially facilitates PAR2 cleavage, has not been addressed. Thus, we examined whether TM-bound thrombin cleaved PAR2 and tested biological responses such as nuclear factor kappa B (NF-κB) DNA binding activity and cytokine release.

**Materials and Methods:** We examined 293T cells overexpressing PAR2 and TM for thrombin recruitment by TM promoting PAR2 cleavage. To test for the TM–thrombin interactions required for PAR2 cleavage and to map cleavage sites on PAR2, mutant constructs of TM or PAR2 were engineered. Biological effects because of PAR2 activation were investigated using an NF-κB reporter system and cytokine release.

**Results and Conclusions:** We identified that, at low to moderate concentrations, thrombin cleaved PAR2 in a TM co-receptor-dependent manner with cleavage efficiency comparable to that of trypsin. In TM’s presence, thrombin efficiently cleaved both, PAR1 and PAR2, albeit kinetics differed. Whereas the majority of surface expressed PAR1 was immediately cleaved off, prolonged exposure to thrombin resulted in few additional cleavage. In contrast, PAR2 cleavage was sustained upon prolonged exposure to thrombin. However, TM EGF-like domain 5 was required and TM chondroitin sulfate (CS) proteoglycan sites serine 490 and serine 492 assisted in PAR2 cleavage, while thrombin preferentially cleaved at arginine 36 on PAR2’s N-terminus. Note that thrombin-induced activation of NF-κB via PAR2 resulted in release of interleukin-8. Thus, we provide a novel concept of how thrombin efficiently cleaves PAR2 in a TM-dependent manner, resulting in pro-inflammatory interleukin-8 release. This unexpected pro-inflammatory role of TM, promoting cleavage and activation of PAR2 by thrombin, may lead to novel therapeutic options for treating inflammatory and malignant diseases.

## Introduction

Clotting proteases elicit multiple physiological and pathophysiological responses beyond clot formation. The family of PARs has evolved to allow cells to react to presence of clotting (serine) proteases [1]. PARs belong to the seven-transmembrane G-protein-coupled receptor family. In mammals, the family of PARs consists of four highly related serine protease receptors, i.e., PAR1–4 [2–7].

Previously, PAR2 has been linked to inflammation-driven diseases of the brain, cardiovascular system, lungs, joints, and gastrointestinal tract [8–16], as well as cancer growth [17, 18], migration, and metastasis [9, 19]. Despite its importance in pathophysiological processes, no treatments based on interference with PAR2 have been established [20]. There are potential obstacles for defining therapeutic targets because of the existence of multiple activation mechanisms and a large variety of causal activating proteases [21–23], as well as the complex nature of PAR2 compartmentalization and co-localization [19]. Recent studies that emphasize the importance of PAR hetero-dimerization [24], requirement of PAR2 co-localized co-receptors [22], and introduction of the concept of biased PAR agonists [25] further demonstrate the complexity of PAR2-related functions under physiological or pathophysiological conditions.

To date, PAR2 has been assumed to resist activation by the prototypical procoagulant protease thrombin, as it has not been observed to activate overexpressed PAR2 in the absence of co-receptors [1, 5, 26]. Moreover, only extremely high, potentially supra-physiological concentrations of thrombin (up to 500 nM) cleave peptides that are homologous to PAR2’s N-terminus and activate PAR2 [27].

Unlike previous observations of PAR2 resisting thrombin activation, *in vitro* and *in vivo* models have demonstrated multiple effects involving both, PAR2 and thrombin, such as cancer cell invasion [18], signal transduction [28, 29], and late sepsis. To explain these observations, PAR1 and PAR2 transactivation is the leading model; the thrombin-generated N-terminus of PAR1 signaling via a PAR2 hetero-dimer partner [24, 30].

To date, the activation of PAR2 by cofactor (or co-receptor)-bound thrombin has not been systematically evaluated, although co-receptor-dependent mechanisms, such as tissue factor-bound clotting factor VIIa activating PAR2 [22], endothelial protein C receptor-bound activated protein C (aPC) cleaving PAR1 and PAR3 [31], endothelial protein C receptor-bound clotting factor Xa cleaving PAR1 [32], and activating PAR1 or PAR3 [33–35], are well known. Moreover, in mice, PAR3 recruits thrombin which facilitates cleavage of PAR4 [36].

For thrombin-mediated activation of protein C, the cell surface glycoprotein TM (also known as fetomodulin or CD141) serves as a cofactor [37,38]. TM is primarily expressed on all vascular endothelial cells [39], keratinocytes [40], astrocytes [41], monocytes and macrophages [42], and lung alveolar epithelial cells [43].

Because TM serves as the key re-director of procoagulant thrombin [44], which limits clot formation and injury to the vascular bed [45], TM might play a role in re-directing PAR activation by thrombin in situations of ongoing thrombin generation such as chronic inflammation. Note that TM-bound thrombin changes substrate specificity of thrombin to anticoagulant and anti-inflammatory signaling via protein C activation. TM expression is linked to regulation of homeostatic and inflammatory responses during injuries [46]. For acute injuries of the endothelium, TM downregulation or shedding occurs; accordingly, soluble TM (sTM) serves as a marker for acute inflammation [47]. Furthermore, compared with the endothelium, less is known about the TM co-receptor’s function in other tissues.

Therefore, we tested whether TM serves as a co-receptor for the thrombin-dependent cleavage of PAR2. Moreover, we verified whether TM-bound thrombin activation of PAR2 resulted in pro-or anti-inflammatory effects.

## Materials and Methods

### Reagents

Thrombin [EC 3.4.21.5] was purchased from Haematologic Technologies (Essex Junction, VT, USA) and trypsin [EC 3.4.21.4] from Gibco (Thermo Fisher Scientific, Reinach, Switzerland). Recombinant aPC (Xigris) was purchased from Eli Lilly and Company (Indianapolis, IN, USA). Peptides corresponding to the N-terminus of R41-cleaved PAR1 (TFLLRNPNDK), R36-cleaved PAR2 (SLIGRL), and PAR4 (AYPGKF) were custom-made (Synpeptide Co., Ltd., Shanghai, China; and ProteoGenix, Schiltigheim, France).

Recombinant sTM (SRP3172), Nordihydroguaiaretic acid (NDGA), TNFα and PI3K inhibitor LY294002 were purchased from Sigma (Buchs SG, Switzerland). We used lepirudin (Refludan; Schering, Berlin, Germany) to block thrombin. In fact, when added without thrombin, lepirudin had no effect in any of the reported assays and readouts. Heparin was purchased from Bichsel (Interlaken, Switzerland). Thrombin exosite I blocking aptamer (5′-GGTTGGTGTGGTTGG-3′ [48]) and exosite II blocking aptamer (5′-AGTCCGTGGTAGGGCAGGTTGGGGTGACT-3′ [49]) were custom-made (Microsynth, Balgach, Switzerland). Monoclonal anti-PAR1 ATAP2 was used as described previously [32, 50] and anti-PAR2 (SAM11; Thermo Fisher Scientific), anti-PAR2_#344222 (MAB3949; R&D Systems, Inc., MN, USA), anti-TM_#1009 [141C01(1009), Thermo Fisher Scientific], anti-TM_#2375 (American Diagnostics, Pfungstadt, Germany), anti-TM_#501733 (MAB3947; R&D Systems), anti-His_#AD1.1.10 (MAB050; R&D Systems), and goat anti-mouse IgG Alexa-594 (R37121; Thermo Fisher Scientific) were all used according to manufacturer’s instructions unless stated otherwise. All oligonucleotides were purchased from Microsynth. For quantifying luciferase activity, the Bright-Glo™ Luciferase Assay System (Promega, Dübendorf, Switzerland) was used, in accordance to the manufacturer’s instructions.

### Peptide cleavage

A high-binding microplate (#9018; Corning, Tewksbury, MA, USA) was coated (o/n) with anti-His#AD1.1.10 (2 µg/ml) and anti-TM_#501733 (1 µg/ml). Then, the plate was washed and incubated with buffer or recombinant sTM (60 nM) to bind to anti-TM for 30 minutes. Synthetic PAR2 peptide (10 µg/ml) was bound to the plate by the N-terminal His-tag (1 h). We added pre-warmed thrombin (30 nM) to the washed plate and incubated the plate for 1 h at 37°C. The uncleaved (full length) PAR2 peptide was detected via the C-terminal biotin-tag using streptavidin-HRP and quantified by a peroxidase-based enzyme immunoassay.

### Cell culture, plasmid transfection, and gene silencing

Epithelial A549 cells and human embryonic kidney cell-derived 293T cells (ATCC; LGC Standards GmbH, Wesel, Germany) were cultivated and propagated as described previously [32, 50]. In brief, cells were propagated in DMEM (Gibco) containing 10% fetal calf serum (GE Healthcare, Glattbrugg, Switzerland), detached using dissociation buffer (Gibco), transfected, seeded, and grown to confluence in multi-well dishes. Two days later, experiments were performed using washed cells and serum-free conditioning medium (DMEM) containing 0.04% BSA (Sigma).

Tagged and untagged PAR1, PAR2, TM constructs, and pGL4.32[*luc2P*/NF-κB-RE/Hygro] (Promega) were transiently expressed in 293T cells by transfection with Lipofectamine2000 (Thermo Fisher Scientific), as described previously [34]. Cells used for cleavage assays were transfected with 1.6 µg/ml plasmid DNA; however, for NF-κB luciferase and IL-8 enzyme-linked immunosorbent assay (ELISA) assays, 1.0 µg/ml plasmid DNA was used for transfection of the 293T cells. To reach comparable amounts of PAR2 expression, in those cells that overexpressed several proteins (such as with and without TM), the empty pcDNA3.1/Zeo(+) plasmid (Thermo Fisher Scientific) was used as a mock plasmid. We performed gene silencing using Lipofectamine RNAiMAX (Thermo Fisher Scientific) as described previously [32], using 20 nM silencing RNA (siRNA). GGAACCAAUAGAUCCUCUAtt (si PAR2) to silence PAR2 and AGGUAGUGUAAUCGCCUUGtt was used as a control silencing RNA (si control). Furthermore, all oligonucleotides were synthetized by Microsynth.

### Overexpression constructs

The overexpression constructs were embedded within the pcDNA3.1/Zeo(+) (Thermo Fisher Scientific) expression plasmid, as described previously [34]. TM was cloned from cDNA (obtained from RNA isolated from EA.hy926 endothelial cells) using primers containing restriction sites (italics) HindIII and XhoI (forward: 5′-GCTA*<u>AAGCTT</u>*GACAGGAGAGGCTGTCGCCATC-3′ and reverse: 5′-GCTA*C<u>TCGAG</u>*GGGAATAAGTGGGGCTTGCT-3′). The HindIII/XhoI-digested PCR product was directly introduced into the HindIII/XhoI-cut pcDNA3.1/Zeo(+) backbone. For removal of N-terminal domains in TM, two unique BamHI restriction sites were introduced by mutagenesis (NEB’s plasmid mutation kit; Ipswich, MA, USA): the first 3′ from the signal sequence and the second 3′ from the sequence that encoded the domain to be removed. Digestion of the mutated TM expression vector with BamHI followed by plasmid re-ligation allowed dropping domains while maintaining the TM native signal sequence (details are provided in Supplementary Table S1). Note that TM chondroitin sulfate (CS) sites serine 490 and 492 were mutated to alanine with the primer set (forward: 5′-CACGGCTCGACCTCAATG-3′, reverse: 5′-GGGGCTCGCC*<u>AGC</u>*GCC*<u>GGC</u>*GTCGCCACCGTCC-3′).

PAR1 and PAR2 alkaline phosphatase-tagged reporter (“AP-PAR1 and AP-PAR2”) was as described elsewhere [34]. Sequence of the PAR4 alkaline phosphatase-tagged reporter (“AP-PAR4”) is described in Supplementary Table S4. In brief, an alkaline phosphatase (AP) tag (pSEAP; Clontech, CA, USA) was linked to the N-terminus of PAR1 cDNA (F2R, NM_001992.4), PAR2 cDNA (F2RL1, NM_005242.5), and PAR4 cDNA (F2RL3, NM_003950.3). Unlike the PAR2 construct mentioned elsewhere [34], an arginine (italics) at the C-terminal end of AP (potential thrombin cleavage site) was removed using phosphorylated mutagenic primers (forward: 5′-CCCGGGTTACTCT*<u>GCG</u>*GCCCAAGGAACCAATAG-3′; reverse: 5′-CTATTGGTTCCTTGGGC*<u>CGC</u>*AGAGTAACCCGGG-3′; Supplementary Table S2). We inserted a green fluorescent C-terminal tag (EGFP; Clontech) analogous to a procedure used in a previous study [34] (for details, see Supplementary Table S2 and Supplementary Scheme S1). To test for preferred cleavage sites on PAR2’s N-terminal, arginine 36 was replaced by alanine (“PAR2 R36A;” protein sequence in Supplementary Table S3) using mutagenic primers (forward: 5′-CCAATAGATCCTCTAAAGGA*<u>GCA</u>*AGCCTTATTGGTAAGG-3′; reverse: 5′-TTCCTTGGGCCGCAGAGTAACCCGGG-3′). Next, all potential thrombin cleavage sites (arginine and lysine) were replaced by alanine (“PAR2 all-to-A” mutant; protein sequence provided in Supplementary Table S3). Note that all constructs that were used were verified by sequencing.

### PAR cleavage reporter assay

The cleavage reporter assay was performed on 293T cells that transiently expressing AP-PAR constructs (Supplementary Table S2), as described elsewhere [32]. In brief, washed cells were incubated with an agonist (for 20 minutes, or if indicated, for repeated periods of 20 minutes), supernatants were removed and filtered (cellulose ester filter; pore size, 0.45 µm), and AP activity was quantified (1-Step PNPP; Thermo Fisher Scientific) by spectrophotometry (Labsystems Multiskan MCC/340; Fisher Scientific). For all experiments, AP-PAR expression levels were confirmed to be comparable among constructs by quantifying AP activity of cell surface. Data are presented in optical density (OD) after subtraction of signals obtained from cells not incubated with the agonist.

### Immunoassays

IL-8 was quantified by a commercial (R&D Systems) sandwich ELISA, according to the manufacturer’s instructions. In brief, 293T cells were transfected with plasmid DNA and A549 cells that were silenced with siRNA 48 h prior to incubation with agonists for 24 and 6 h, respectively. Then, IL-8 release was measured in supernatants. As described previously, cell surface PAR2 and TM were quantified by cell surface ELISA [50].

### Immunofluorescence microscopy

EGFP-tagged protein (Supplementary Scheme S1) was overexpressed using the same protocols as that described for PAR cleavage reporter constructs and visualized using appropriate filter sets of an Axiovert-10 fluorescent microscope (Carl Zeiss AG, Feldbach, Switzerland). As described for the cell surface ELISA, untagged proteins were incubated with antibodies; however, the secondary antibody carried an Alexa-594 fluorescent tag (Thermo Fisher Scientific) rather than HRP.

### NF-κB luciferase assay

293T cells were transfected with the NF-κB firefly luciferase construct pGL4.32[*luc2P*/NF-κB-RE/Hygro] (Promega) along with AP-PAR constructs and seeded into black, clear-bottomed 96-well plates (Greiner Bio-One, Kremsmünster, Austria). After transfection for 48 h, cells were washed and incubated with an agonist (for 6 h) at 37°C. Bright-Glo™ Luciferase Assay System reagent was added to cells and luciferase activity was detected via luminescence (SpectraMax i3; Molecular Devices, San Jose, CA, USA), according to the manufacturer’s instructions. Furthermore luciferase activity was normalized to the buffer-induced relative light units (RLUs).

### Statistics

We analyzed and presented the data using GraphPad Prism5 (GraphPad Software, La Jolla, CA, USA). To calculate the indicated P-values, a two-sample, two-tailed homoscedastic t-test and one-way or two-way ANOVA with Bonferroni correction were used.

## Results

### Effect of PAR2 activation by thrombin on IL-8 release in native A549 cells

Various studies showed expression of all PARs in lung epithelial cells promoting IL-8 release upon PAR agonist peptide stimulation due to PAR cis- or trans-activation [51] [52]. In this study, we first tested the concentrations of PAR agonist peptides required to induce IL-8 secretion in A549 cells. Similar to a previous study [53], we found that A549 cells were induced to release IL-8 on incubation with the PAR2 agonist peptide SLIGRL, whereas the PAR1 agonist peptide TFLLRNPNDK and the specific PAR4 agonist peptide AYPGKF demonstrated no such effect, even at high concentrations. Unlike the PAR1-specific peptide, thrombin significantly induced IL-8 release from 30 to 100 nM (Figure 1A). To test whether thrombin induced IL-8 release could be linked to PAR2 activation, we relied on the PAR2-specific inhibitor NDGA. NDGA did neither affect IL-8 release at baseline (buffer control) nor in highly TNFα-induced A549 cells (not shown). However, NDGA significantly reduced IL-8 release by thrombin as well as by the PAR2-specific agonist SLIGRL (Figure 1B). Consistently, silencing of PAR2 by siRNA abolished responses of A549 cells to thrombin or SLIGRL induction (Figure 1C). In summary, IL-8 release by thrombin stimulation was significantly reduced in PAR2-silenced or inhibited lung epithelial cells, suggesting that thrombin plays a role in PAR2 signaling.

**Figure 1:**
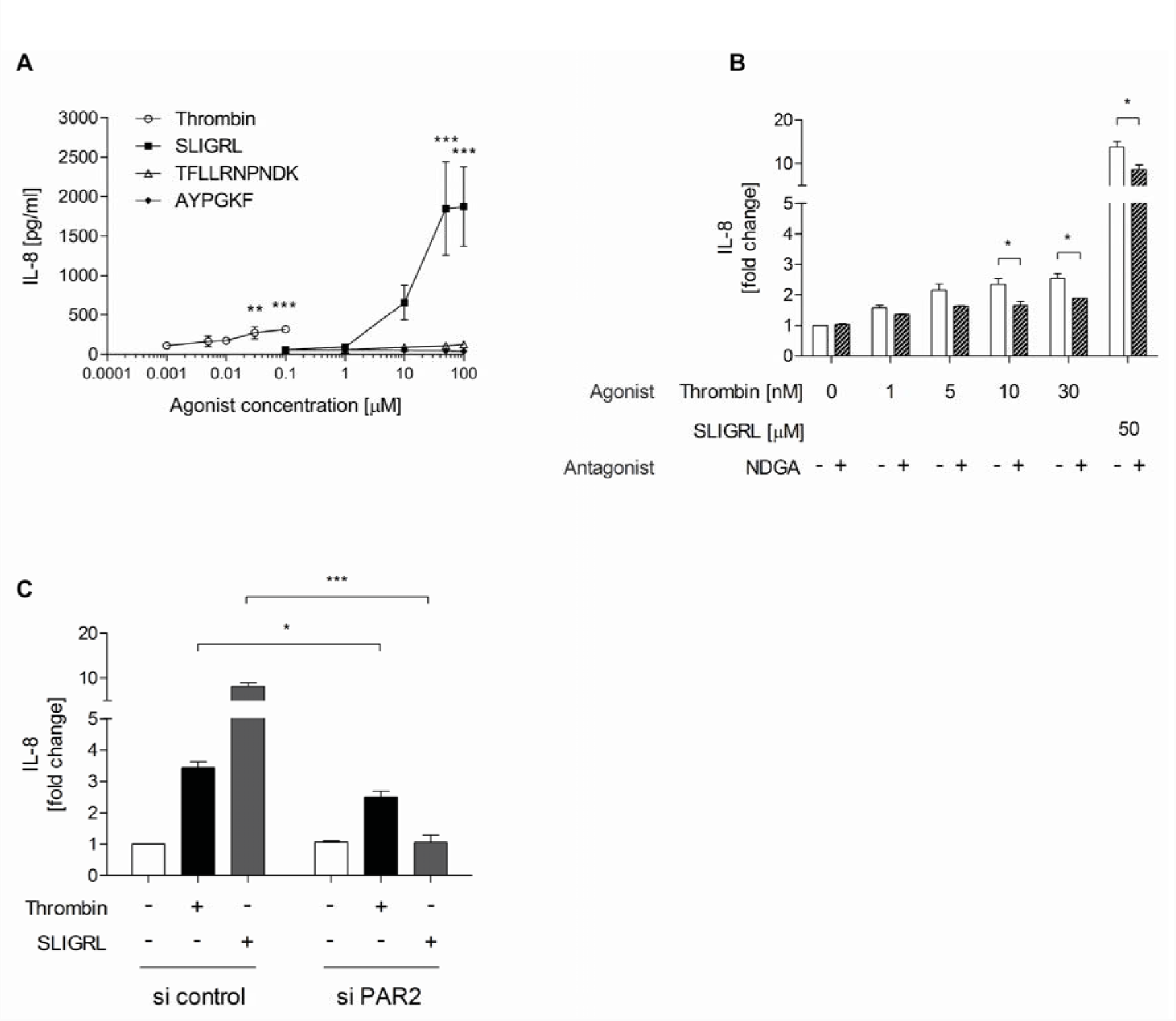
Increased IL-8 release upon thrombin and SLIGRL stimulation in a PAR2-dependent manner. (**A**) A549 cells were stimulated with thrombin, PAR2 agonist peptide SLIGRL, PAR1 agonist peptide TFLLRNPNDK, and PAR4 agonist peptide AYPGKF for 6 h. Then, IL-8 release was quantified by a peroxidase-based enzyme immunoassay. (**B**) A549 were pretreated with DMSO or NDGA (20 µM) for 30 min before stimulation with buffer, thrombin or SLIGRL (concentrations indicated in the graph). IL-8 release in the supernatant was quantified by ELISA after 6 h of agonist incubation. (**C**) A549 cells were transfected with siRNA targeting no specific mRNA (si control) or siRNA targeting PAR2 (si PAR2). Then, cells were stimulated with buffer, thrombin (30 nM), or SLIGRL (50 µM) for 6 h before IL-8 release was measured in cell supernatants. Data was presented as mean ± SEM; three independent experiments were each performed in triplicate; ns *p* > 0.05, * *p* < 0.05, ** *p* < 0.01, *** *p* < 0.001, **** *p* < 0.0001 using one-way ANOVA (A) and student’s *t* test (B,C).

### Role of TM on PAR2 cleavage by thrombin

To test whether IL-8 induction by thrombin in epithelial cells of the lung could be caused by direct cleavage of PAR2 using a co-receptor, we established a purified system comprising immobilized PAR2 peptide with immobilized sTM and thrombin. In this purified system, thrombin efficiently cleaved the PAR2 peptide in immobilized sTM’s presence; however, consistent with the established conclusion, thrombin failed to directly cleave PAR2 in the co-receptor’s absence (Figure 2A). To test on human cells whether physiologically achievable concentrations of thrombin cleaved PAR2 and whether co-receptor(s) could be involved, we relied on the previously established PAR cleavage reporter system [32, 34, 54]. To allow verification of construct expression by assessing the tag’s cell surface AP activity and monitor cleavage by quantifying the activity of AP released into the supernatant, we overexpressed PAR2 containing an AP enzyme domain at the N-terminus (Supplementary Table S2; Supplementary Scheme S1, [34]). PAR2 reporter and TM constructs were expressed at the cell surface, which was further assessed by fluorescence microscopy and cell surface ELISA. Co-expression of PAR2 and TM did not affect expression of both constructs (Supplementary Fig. S1a and S1b). In such an overexpression system, free thrombin at concentrations up to 30 nM failed to cleave PAR2 (Figure 2B), unlike the results of a recent study [27]. However, in TM’s presence, cleavage efficiency was significantly enhanced and, at low nanomolar concentrations of proteases, thrombin and trypsin showed comparable cleavage efficiency (Figure 2C). To test whether thrombin/TM could mediate release or activation of another protease, which ultimately cleaved PAR2, we added thrombin-induced supernatant from TM-expressing cells to 1) PAR2- and 2) PAR2- and TM-expressing cells, for assessing PAR2 cleavage. We identified that, in the absence of TM, PAR2 was not cleaved and, wherever thrombin in supernatant was blocked by the specific inhibitor lepirudin, cleavage was blunted (Figure 2D). Addition of activated protein C or inhibition of metalloproteases did not affect PAR2 cleavage by thrombin (Supplementary Fig. S3a and S3b). Therefore, overall, these results cannot support a model whereby thrombin induces a PAR2-cleaving protease. To simulate more physiological conditions where PAR1, PAR2 and TM are co-expressed, and to test whether the co-expression impacts cleavage, we overexpressed 1) PAR1 containing an N-terminal AP enzyme tag together with untagged-PAR2 or 2) PAR2 containing an N-terminal AP enzyme tag together with an untagged-PAR1 (Supplementary Table S2; Supplementary Scheme S1, [34]) in presence or absence of TM. As expected, PAR1 was rapidly cleaved by thrombin (5 and 30 nM) while PAR2 - in TM’s presence - was more efficiently cleaved upon prolonged thrombin exposure (5 and 30 nM) (Figure 2E and 2F). The cleavage of PAR2 at 60 minutes of exposure to thrombin (30 nM) was comparably efficient to the PAR1 cleavage at 20 minutes (Figure 2F). Cleavage of PAR2 was sustained upon prolonged exposure to thrombin for up to 25 hours (Supplementary Fig. S4a-c). Note that, upon a exposure to thrombin for 3 hours, a broad range of thrombin concentrations from 1 to 30 nM resulted in significant cleavage of PAR2 (Supplementary Fig. S4a-c). Our observations suggest that TM-bound thrombin directly cleaves PAR2 in our overexpression system and that TM dependent cleavage of PAR2 by thrombin is enhanced over time. Thus, in summary, our data suggest that thrombin at moderate concentrations (5 and 30 nM) efficiently cleaves overexpressed PAR2 reporter construct in TM’s presence.

**Figure 2:**
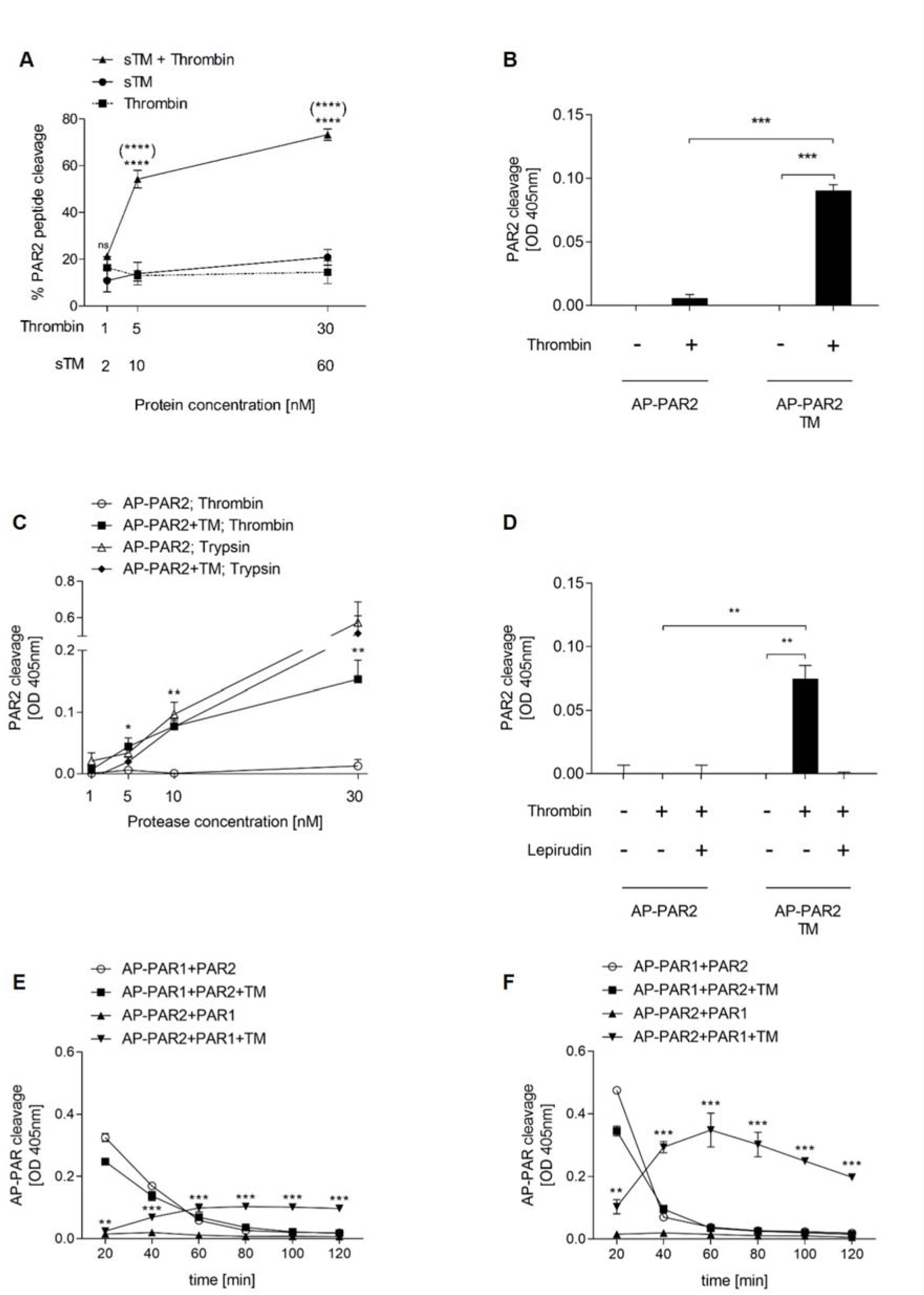
Thrombin efficiently cleaves overexpressed PAR2 in the presence of TM. (**A**) A PAR2-derived synthetic peptide mimicking the entire extracellular N-terminal domain of PAR2 was bound to the plate via a C-terminal 6xHis-tag and incubated with thrombin in the presence or absence of immobilized sTM. Uncleaved PAR2 was detected via ELISA peroxidase-based quantification of unremoved N-terminal biotin tags. Stars indicate comparisons of **\*** PAR2 peptide cleavage of sTM+thrombin vs. thrombin without sTM or (**\***) sTM+thrombin vs. sTM without thrombin. (**B**) 293T cells transiently overexpressing AP-PAR2 reporter construct either alone or with TM. Where indicated, cells were incubated with thrombin (30 nM) for 20 minutes before the released AP’s activity was measured in the cells’ supernatants, serving as a surrogate for PAR2 cleavage. (**C**) Cells expressing TM and AP-PAR2 reporter construct were incubated with agonists as indicated and assayed as described in (B). Stars indicate the comparison of thrombin to buffer in cells overexpressing AP-PAR2+TM. (**D**) 293T cells expressing only TM were incubated with thrombin (20 min, 30 nM). Thereafter, the supernatant was swapped to AP-PAR2- or AP-PAR2 and TM-expressing cells and the cells were incubated for another 20 minutes. Where indicated, the supernatant was pre-incubated with the thrombin inhibitor lepirudin (30 nM) before it was added to the cells expressing AP-PAR2 or AP-PAR2 and TM. (**E**) and (**F**) 293T cells expressing AP-PAR1 together with non-tagged PAR2 or AP-PAR2 together with non-tagged PAR1 with or without TM were incubated with (**E**) thrombin (5 nM) or (**F**) thrombin (30 nM) for 20 minutes. AP activity was measured in the supernatant and the cells were agonist-incubated for repeated periods of 20 minutes. Stars indicate the comparison of thrombin to buffer in cells overexpressing AP-PAR2+PAR1+TM. Data are presented as mean ± SEM; (A-D) three or (E,F) four independent experiments, each performed in triplicate; ns *p* > 0.05, * *p* < 0.05, ** *p* < 0.01, *** *p* < 0.001, using student’s *t* test.

### Importance of interactions between TM EGF-like domains and CS with thrombin exosite I and exosite II on PAR2 cleavage

Next, we addressed domains of TM required for increasing PAR2 cleavage by thrombin. TM contains six epidermal growth factor (EGF)-like domains, from which EGF-like domain 4 was linked to protein C substrate acceptance, whereas EGF-like domains 5 and 6 were linked to thrombin recruitment [55] to the cell surface. To test whether these EGF-like domains were involved in promoting PAR2 cleavage, we produced N-terminally truncated mutants of TM (Supplementary Table S1). The mutants were expressed and recognized as expected by cell surface ELISA using anti-TM targeting EGF-like domains 5 and 6 (Supplementary Fig. S5). Note that PAR2 was efficiently cleaved when wild-type TM or mutants containing EGF-like domains 5 and 6 were present. The TM mutant composed of only EGF-like domain 6 failed to promote the cleavage of PAR2 by thrombin (Figure 3A).

**Figure 3:**
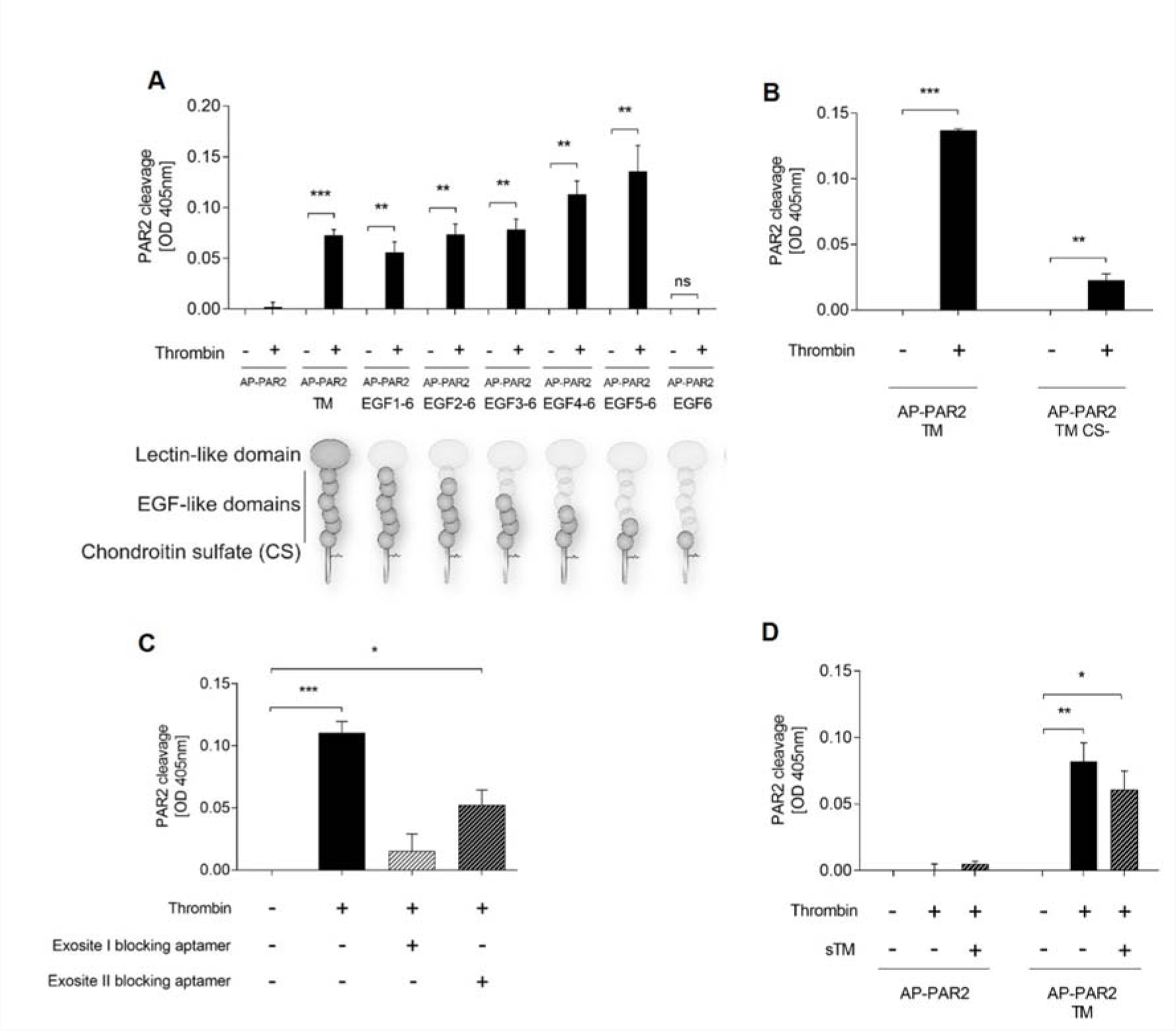
For PAR2 cleavage by thrombin, interaction between thrombin exosite I and TM’s EGF-like domain 5 is necessary, along with interaction between exosite II and TM’s CS motif. (**A**) 293T cells overexpressing AP-PAR2 reporter construct alone or together with either wild-type TM (TM) or mutants of TM lacking N-terminal domain(s). Scheme of the expressed TM domains is provided (bottom of A) and the details are explained in Supplementary Table S1. After 20 minutes of incubation with thrombin (30 nM), PAR2 cleavage was measured. (**B**) AP-PAR2 reporter construct was co-expressed in 293T cells with wild-type TM or a TM mutant lacking the CS sites serine 490 and 492 (TM CS-). Moreover, PAR2 cleavage by thrombin (30 nM) was measured as in (A). (**C**) 293T cells overexpressing AP-PAR2 and TM were incubated (for 20 minutes) with buffer, thrombin (30 nM), or a mixture of thrombin (30 nM) and exosite blocking aptamers (2 µM); then, PAR2 cleavage efficiency was measured. (**D**) 293T cells overexpressed AP-PAR2 with or without TM. Where indicated, cells were incubated for 20 minutes with thrombin (30 nM) alone or a 1:1 mixture of thrombin and sTM (20 minutes of pre-incubation at 37°C for complex formation). Data are presented as mean ± SEM; three independent experiments were each performed in triplicate; ns *p* > 0.05, * *p* < 0.05, ** *p* < 0.01, *** *p* < 0.001, using student’s *t* test.

Moreover, interactions between thrombin exosite II and CS on TM were involved for thrombin recruitment [56] and to enhance the endothelial protein C pathway. To test whether such interactions between exosite II and CS facilitated PAR2 cleavage by thrombin, we synthesized a TM mutant that lacked the serine 490 and 492 glycosylation sites. Similar to the conclusions from the protein C pathway that interactions between exosite II and CS assist with thrombin recruitment toward TM, CS-deficient mutant of TM was less efficient for supporting the cleavage of PAR2 by thrombin (Figure 3B). Similarly, blocking the interactions of exosite II with CS with an aptamer, a short single-stranded DNA specifically blocking exosite II [49], interfered with PAR2 cleavage (Figure 3C). Supporting this observation, PAR2 cleavage was reduced by exosite II binding agent heparin (Supplementary Fig. S2). Unlike exosite II and TM interaction, the interaction of exosite I with EGF-like domain 5 and 6 of TM is considered essential for thrombin recruitment [57]. Moreover, blocking exosite I by an aptamer [48] blunted PAR2 cleavage by thrombin (Figure 3C).

To test whether sTM supports PAR2 cleavage by thrombin, thrombin was pre-incubated with recombinant sTM that lacks the membrane anchor. The mixture was then tested for PAR2 cleavage. In the absence of cell surface-expressed TM, sTM– thrombin mixture failed to significantly cleave PAR2 (Figure 3D). This observation supports either the lack of sTM’s role in assisting PAR2 cleavage by thrombin or the failure of our cells to recruit sTM–thrombin complexes onto the cell surface.

Thus, similar to our observations of the protein C pathway, our data suggest that, for efficient PAR2 cleavage by thrombin, interactions between TM’s EGF-like domain 5 and the thrombin exosite I is particularly important, in addition to the efficiency-boosting effect of the interaction between CS and thrombin exosite II, as well as the essential role played by cell surface-anchored TM.

### Role of PAR2 N-terminal arginine 36 in cleavage by thrombin

Previous evidence suggested that PARs harbor a unique cleavage site that promotes initiation of biological effects [1]. Recently, however, studies have revealed that PARs behave as “multi-switches” with several cleavage sites, resulting in cleavage-site-specific biological effects [23, 34, 35, 58] and referred to as biased signaling. Thus, we investigated whether TM-bound thrombin cleaved PAR2 at specific sites. Similar to our studies in PAR1 and PAR3 [34], we synthesized mutants of PAR2 with alanine (A) substitutions for the positively charged amino acids arginine (R) and lysine (K) in the extracellular N-terminal domain (Supplementary Table S3). A mutant of PAR2 that was devoid of all R and K on the entire N-terminal domain (“PAR2 all-to-A” mutant; Supplementary Table S3) resisted cleavage by thrombin (Figure 4A). Removal of canonical R36 (“PAR2 R36A” mutant; Supplementary Table S3) yielded an almost thrombin-resistant mutant (Figure 4B); however, additional sites at which cleavage occurs inefficiently might be present. Thus, our data suggest that R36 is the preferred cleavage site of TM-bound thrombin; however, the existence of additional cleavage site(s) cannot be ruled out.

**Figure 4:**
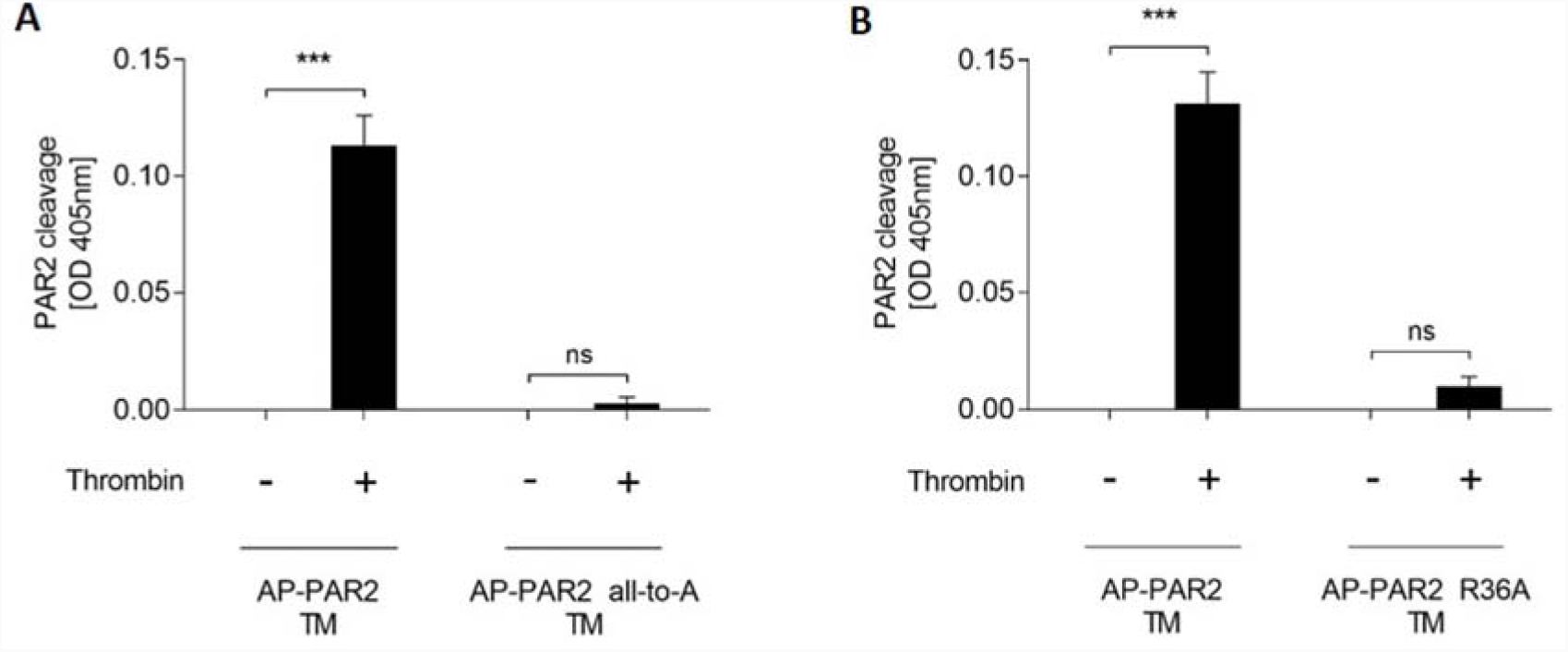
TM-bound thrombin preferentially cleaves PAR2 at arginine 36. (**A**) and (**B**) 293T cells overexpressing TM and wild-type or mutants of AP-PAR2 reporter constructs. The mutant “AP-PAR2 all-to-A” contained no positively charged amino acids in the N-terminal PAR2 domain and in the “AP-PAR2 R36A” mutant, arginine 36, was replaced by alanine (details provided in Supplementary Table S3). Cells were incubated with thrombin (30 nM) for 20 minutes before PAR2 cleavage was measured. Data are presented as mean ± SEM; three independent experiments were each performed in triplicate; (A-B) ns *p* > 0.05, * *p* < 0.05, ** *p* < 0.01, *** *p* < 0.001, using student’s *t* test.

### Role of the NF-κB pathway downstream of thrombin-cleaved PAR2

Previously, pro-inflammatory signaling was reported to occur because of PAR2 activation by other proteases that cleaved PAR2 at R36 [11–13]. Thus, we investigated whether TM-bound thrombin cleaved PAR2 and induced pro-inflammatory signaling via NF-κB activation. In our commercial NF-κB reporter system, thrombin failed to induce NF-κB activation when PAR2 and TM were absent or expressed alone (Supplementary Fig. S6a). NF-κB DNA binding activity, however, increased in a manner dependent on thrombin concentration due to the availability of PAR2 and TM (Figure 5A). PAR2 agonist peptide SLIGRL induced NF-κB DNA binding activity in PAR2 and TM overexpressing cells similarly to thrombin; however, >1000 times higher concentrations of agonist peptide was required to obtain comparable induction (Figure 5A). Similar to thrombin, the PAR1 agonist peptide TFLLRNPNDK induced NF-κB activation only if PAR1 was overexpressed. Similarly, the PAR4 agonist peptide AYPGKF induced NF-κB activation only in cells overexpressing PAR4. As expected, the PAR4 activation by thrombin was less efficient compared to PAR1 activation. In the absence of overexpressed PAR1 or PAR4, agonist peptides showed no such effect, similar to TFLLRNPNDK and AYPGKF failing to activate endogenous PAR1 or PAR4 in these cells and consistent with TFLLRNPNDK and AYPGKF failing to activate overexpressed PAR2 in this system. However, similar to the PAR2 agonist peptide SLIGRL, thrombin-induced NF-κB activation via PAR2 only in the presence of TM (Supplementary Fig. S6b).

**Figure 5:**
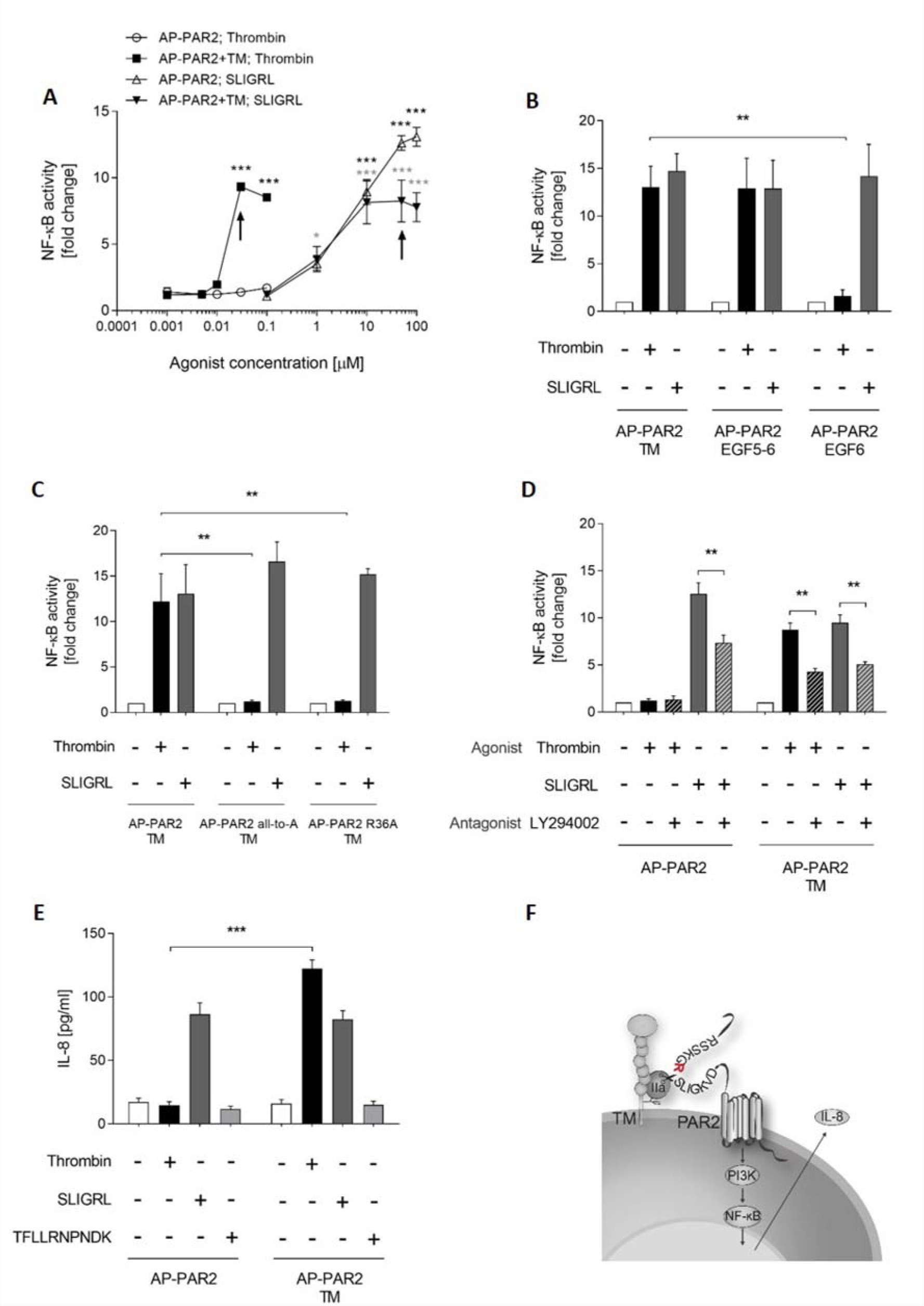
NF-κB DNA binding activity and IL-8 release is induced by TM-bound thrombin in a PAR2-dependent manner. (**A**) 293T cells carrying an NF-κB reporter overexpressed either AP-PAR2 alone or AP-PAR2 together with TM, were incubated with an agonist as indicated and quantified for NF-κB luciferase activity by measuring luminescence. Arrows indicate thrombin (30 nM) and SLIGRL (50 µM), concentrations used in further NF-κB reporter assays. (**B**) Cells overexpressing NF-κB reporter, AP-PAR2 together with wild-type TM, a mutant of TM lacking all domains at the N-terminal side of EGF-like domain 5 (“TM EGF 5–6”), or a mutant lacking all domains at the N-terminal side of EGF-like domain 6 (“TM EGF 6”). Induction of NF-κB luciferase activity in response to the buffer, thrombin (30 nM), and SLIGRL (50 µM) was also measured. (**C**) Cells overexpressing NF-κB reporter, TM, and either AP-PAR2 wild-type or AP-PAR2 mutants that lacked all positively charged N-terminal amino acids (“AP-PAR2 all-to-A”) or only arginine 36 (“AP-PAR2 R36A”). Cells were incubated with buffer, thrombin (30 nM), and SLIGRL (50 µM), after which induction via luciferase was measured. (**D**) Cells overexpressing NF-κB reporter, TM, and AP-PAR2 were pre-incubated with vehicle (DMSO) or the PI3K inhibitor LY294002 (Sigma) for 2 h, followed by incubation with buffer, thrombin (30 nM), or SLIGRL (50 µM) for 6 h before NF-κB luciferase induction was measured. (**E**) 293T cells expressing AP-PAR2 or AP-PAR2 and TM were incubated with an agonist for 24 h, followed by quantification of the release of IL-8 into the supernatant. (**F**) Scheme of proposed PAR2 activation pathway: thrombin (IIa) binds to TM, cleaves PAR2, and activates the NF-κB pathway via PI3K activation. NF-κB activation induces IL-8 synthesis and release. Data are presented as fold induction of buffer RLU, presented as mean ± SEM; three independent experiments were each performed in triplicate; * *p* < 0.05, ** *p* < 0.01, *** *p* < 0.001, using two-way ANOVA (B, C, E), one-way ANOVA (A, D).

Moreover, consistent with our conclusions from the PAR2 cleavage reporter assay, EGF-like domain 5 of TM was required for thrombin- and PAR2-mediated induction of NF-κB activation (Figure 5B).

We then tested whether thrombin’s preferred cleavage site R36 was important for NF-κB induction. In R36’s absence, our assays showed diminished cleavage of PAR2 and no induction of NF-κB activation (Figure 5C).

We then addressed the signal pathways downstream of PAR2 but upstream of NF-κB activation. Phosphatidylinositol 3-kinase (PI3K) pathway is an important regulator of NF-κB activation and potentially links PAR2 cleavage and activation to the NF-κB pathway. To test whether the induction of NF-κB activation after PAR2 cleavage occurred via PI3K, we used the PI3K inhibitor LY294002. Consistent with PI3K linking PAR2 and NF-κB, LY294002 interfered with NF-κB activation by thrombin and activation by PAR2-specific peptide SLIGRL (Figure 5D).

Note that 293T cells secreted IL-8 with the overexpression of PAR2 and SLIGRL agonist induction. However, as shown previously [34] and in Supplementary Fig. S6b, cells remained quiescent upon incubation with the PAR1 agonist peptide, which confirms our previous conclusion that, without transfection, these cells would not respond to PAR agonists. Note that thrombin-stimulated 293T cells overexpressing PAR2 alone remained quiescent. However, in cells expressing TM and PAR2, thrombin (30 nM) and SLIGRL (50 µM) were comparably efficient for inducing the release of IL-8 (Figure 5E). Thus, our data suggest that PAR2 cleaved at R36 by TM-bound thrombin induced NF-κB DNA binding activity via PI3K and ultimately released IL-8 release (Figure 5F).

## Discussion

Our data show, for the first time, efficient TM-dependent direct cleavage and activation of PAR2 by thrombin. This breaks the dogma of PAR2 as the only PAR that resists activation by an important clotting protease thrombin. In a manner dependent on TM co-receptor availability, efficient PAR2 cleavage occurred even at lower concentrations of thrombin. Unexpectedly, at low concentrations of proteases, efficiency levels of PAR2 reporter cleavage between thrombin and trypsin were comparable. PAR2 cleavage by thrombin was - in TM’s presence of TM - comparable to cleavage of PAR1, although kinetics differed. While PAR1 was rapidly cleaved off with minimal sustained cleavage, PAR2 cleavage increased over time. Thus, we suggest that PAR1 is responsible for rapid thrombin responses while PAR2 sustains the thrombin signaling following PAR1 desensitization. In fact, PAR2 cleavage by thrombin was most efficient at the canonical cleavage site arginine 36. Note that this study establishes a direct link of cleavage of PAR2 by thrombin to pro-inflammatory signaling such as activation of NF-κB pathway and release of IL-8 in natively PAR2- and TM-expressing A549 lung epithelial cells. To date, PAR2 has been assumed to be distinct in the family of PARs in terms of resisting cleavage and activation by thrombin [1, 5, 26]. However, biological effects of thrombin have now been linked to PAR2, although the exact mechanism of activation needs to be clarified. Transactivation of PAR2 by thrombin-cleaved PAR1 is one such standard model [24, 59], although it is inconsistent with observations demonstrating that thrombin maintained signaling via PAR2 under conditions where PAR1’s tethered ligand was blocked [9].

As an alternative model, two studies have confirmed extremely high concentrations of free thrombin directly cleaving PAR2 [27, 60]. During a thrombin burst, the required concentration of thrombin of 100–500 nM is potentially achievable [61]. However, in-depth studies in mice that lack the thrombin platelet surface recruiting receptor PAR3 (a nonfunctional receptor in mice) bleed, not in line with the physiologic occurrence of thrombin concentrations at some hundred nanomolar [62]. Studies involving blocking antibodies [63] support an upper limit of physiologically relevant thrombin concentrations of <30 nM, thus ruling out the physiological relevance of PAR2 activation at high concentrations of free thrombin.

Similar to our co-receptor-dependent activation of PARs, as shown in the case of PAR1 by aPC [31] and clotting factor Xa [32], PAR2 by clotting factor FVIIa [22], or platelet-expressed PAR4 by thrombin [62], we postulated that, for efficient PAR2 activation by thrombin, a co-receptor was required. Indeed, we discovered a novel co-receptor function of TM, supporting the cleavage and activation of PAR2 by thrombin. Indeed, TM’s presence increased thrombin’s efficiency for cleaving PAR2. Other potential PAR2 cleaving proteases induced or activated by thrombin or the thrombin-TM complex were ruled out. Highlighting the plausibility of the presence of TM playing a physiological role, 5 nM thrombin and 5 nM trypsin cleaved PAR2 with comparable efficiency. Because TM is not the only cell surface receptor for thrombin [64], other surface recruiting receptors might contribute to further PAR2 activation by thrombin and explain the PAR2-dependent effects of thrombin in cells that did not express TM. We identified that the EGF-like domain 5 of TM and exosite I in thrombin were important for TM-promoted PAR2 cleavage and NF-κB pathway activation by thrombin. Furthermore, this conclusion was supported by removing the EGF-like domains on TM and by blocking thrombin exosite I site using a small specific aptamer. Our observations reinforce previous research on TM [55], showing that in the context of the protein C pathway the described interaction is required for cell surface recruitment of thrombin. Similarly, interactions between exosite II and CS help in thrombin recruitment [57]. Furthermore, we found that exosite II blocking aptamers or exosite II binding heparin interfere with PAR2 cleavage by thrombin. These results are in accordance with a recently published study that links the TM glycosaminoglycan domain to immune responses [65].

Thus, our data confirms that thrombin activates NF-κB and induces IL-8 release for overexpressing 293T cells and triggers IL-8 release in A549 cells in a TM- and PAR2-dependent manner. The specific PAR2 agonist SLIGRL induced IL-8 release, supporting PAR2 stimulation to be sufficient for IL-8 release. Complementary, interfering with PAR2 either via inhibition (NDGA) or knock down (siRNA) significantly reduced responsiveness to thrombin and SLIGRL of natively PAR2 expressing A549 cells. Although thrombin-induced IL-8 release was significantly reduced in PAR2 silenced or inhibited cells, we failed to render our system fully thrombin irresponsive. Possible explanations include incomplete PAR2 knock down, or another (yet to be identified) thrombin receptor induced IL-8 release.

With such a well-characterized system, we provided evidence that co-receptor-bound thrombin directly activates pro-inflammatory pathways via PAR2. Our evidence confirms that A549 cells did not release IL-8 on stimulation with PAR1 agonist peptide. This result is consistent with that of a recent study [53]. Our results support that prolonged thrombin signaling via PAR2 / TM could explain pro-inflammatory signaling in A549 alveolar cells. In view of thrombin inhibition ameliorating chronic inflammation of the lung as reviewed in [66] one is tempted to speculate, that thrombin / TM / PAR2 might serve as underlying mechanism. Thus, stimulation of PAR2 by extravascular thrombin might contribute to inflammatory diseases of the lung.

Our conclusion that thrombin activates PAR2 via the co-receptor TM is heavily reliant on overexpression systems; thus, the physiological relevance of this has yet to be established completely. The confirmation of our concept in a single cell line, i.e., A549 natively expressing PAR2 and TM, cannot eliminate this important limitation. However, to summarize, we provide the first evidence that TM-bound thrombin cleaves and activates PAR2 efficiently at low to moderate nanomolar concentrations and that this pathway of NF-κB activation results in the release of IL-8 in PAR2- and TM-overexpressing and natively expressing cells. Further studies will be required to link this novel signaling pathway to important pathophysiological processes, particularly for chronic inflammatory diseases such as asthma, arthritis, and cancer. Note that such studies may reveal novel therapeutic strategies for diseases that are linked to PAR2.

## Acknowledgments

We would like to express our sincere gratitude for the support given to RAS by the Vontobel Stiftung, the Swiss National Science Foundation grant #PZ00P3_136639, the Hartmann Müller-Stiftung, the University Hospital Zurich, and the University of Zurich.

## Authors’ Contributions

Study designed by DMH and RAS. Experiments conducted by DMH, AGF, and JM. Data analysis and interpretation by DMH and RAS. Manuscript preparation by DMH and RAS. All authors read and approved the final manuscript.

## Disclosure of Conflict of Interest

The authors state that they have no conflicts of interest.

## Abbreviations

PAR: protease-activated receptor
TM: thrombomodulin
CS: chondroitin sulfate
EGF-like domain: epidermal growth factor-like domain

