## Supplementary data for "Thrombin cleaves and activates the protease-activated receptor 2 dependent on thrombomodulin co-receptor availability"

**Supplementary Fig. S1a**

**Supplementary Fig. S1b**

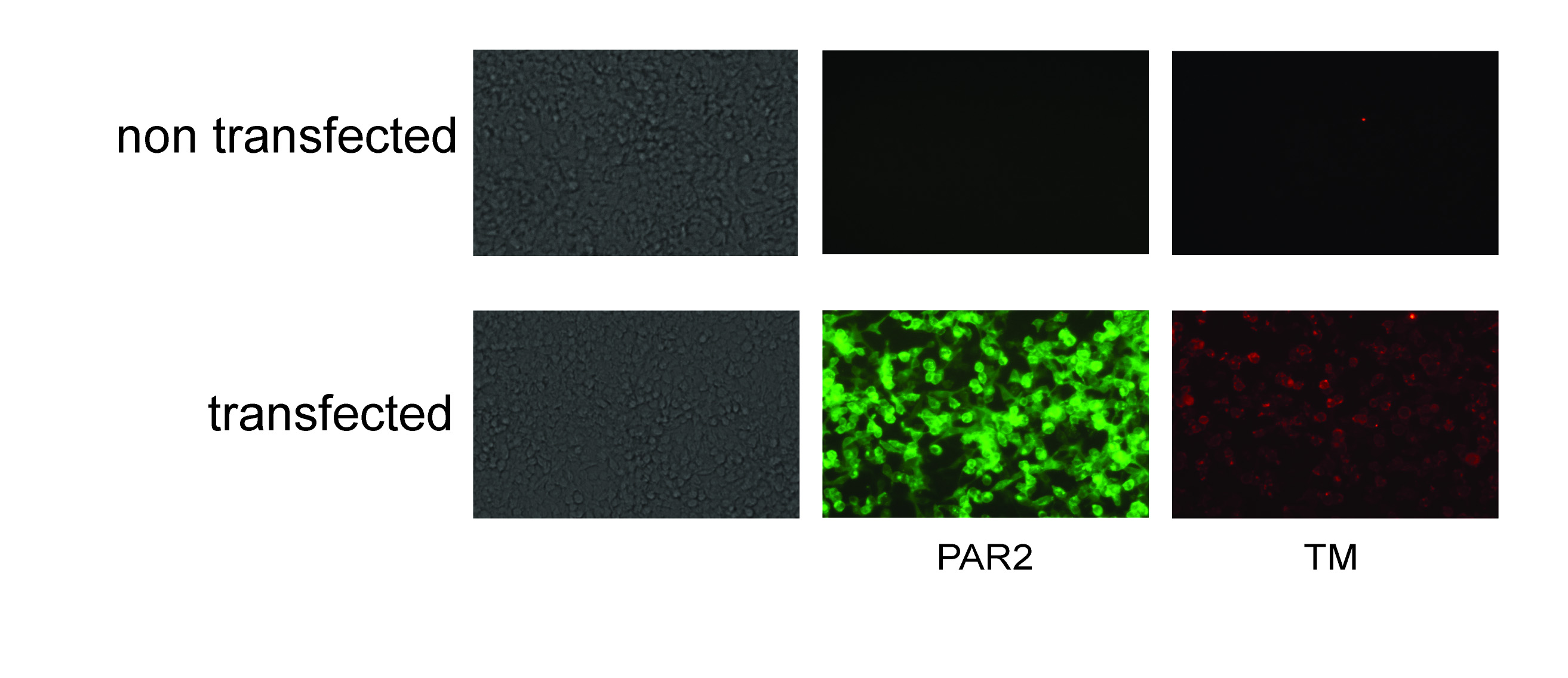

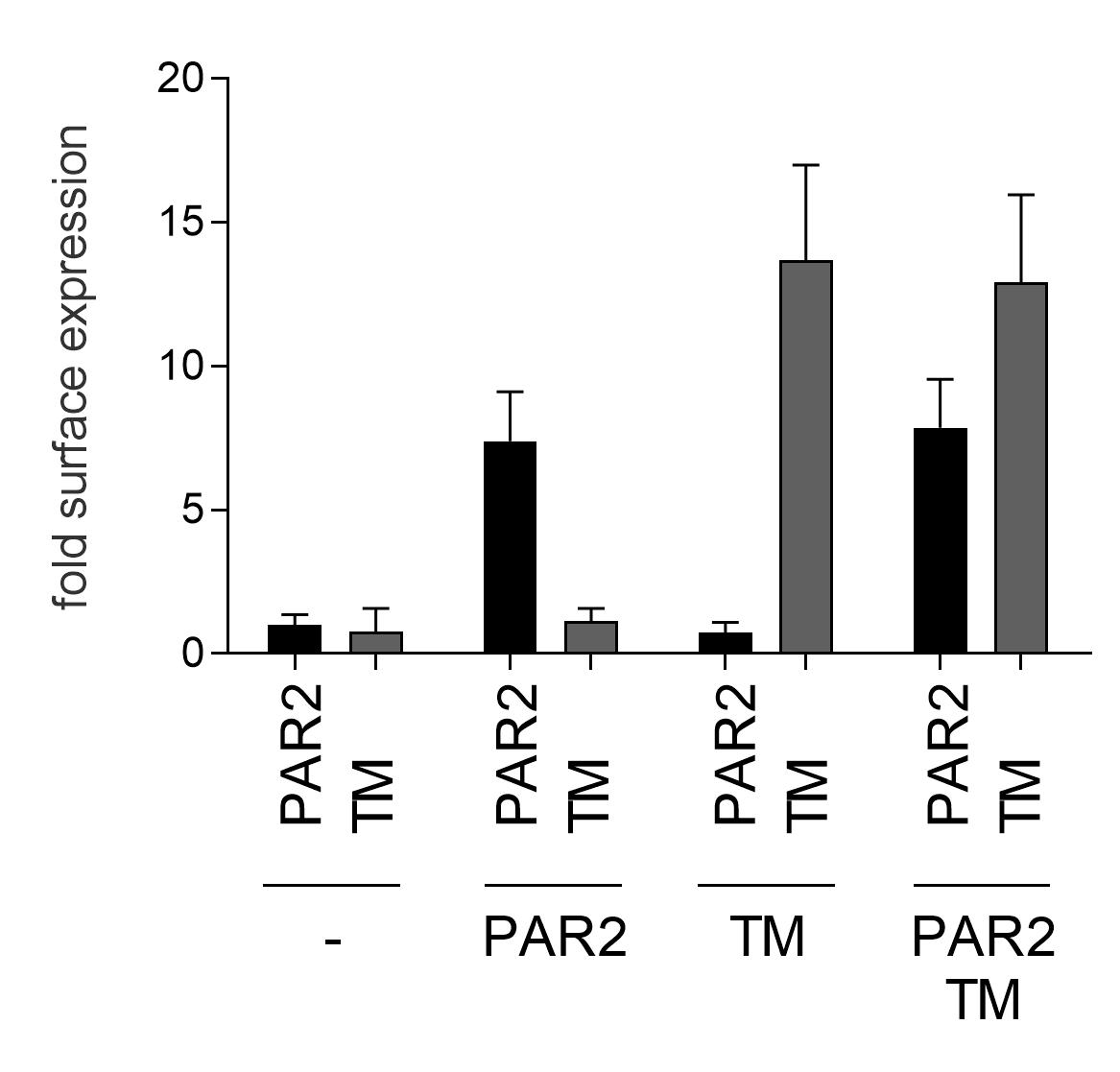

**Supplementary Fig. S1: Visualization and quantification of overexpressed PAR2 and TM.** (**Fig.** **S1a**) Representative microscopic pictures (magnification 10Х) either native 293T cells (upper line) or cells transiently expressing TM together with a PAR2 construct carrying a C-terminal EGFP tag (Supplement Table S2; Supplement Scheme S1). The EGFP tag allowed direct visualization of PAR2 while TM was stained with anti-TM (American Diagnostics #2375) followed by goat anti-mouse Alexa 594 and visualized (dark field, 1^st^ column; fluorescence at 450nm, 2^nd^ column; 620nm 3^d^ column). (**Fig. S1b**) Cell surface ELISA of 293T cells overexpressing AP-PAR2 or TM alone or AP-PAR2 and TM together. Cells were PFA fixed, stained with anti-PAR2#344222 (MAB3949; R&D Systems, Inc. 1:100) or anti-TM#1009 (141C01(1009)), Thermo Fisher Scientific;1:100) followed by an HRP-coupled anti-mouse and photometric quantification of the specific signal at OD 450-690nm. Data presented as mean +/- SEM; 3 independent experiments each performed in triplicate.

**Supplementary Fig. S2**

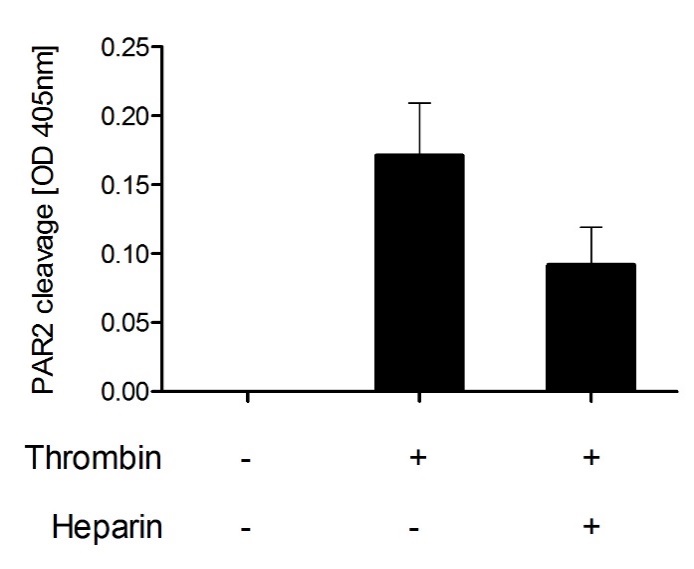

**Supplementary Fig. S2: PAR2 cleavage assay with the thrombin inhibitor heparin.** Cleavage assay of 293T cells overexpressed AP-PAR2 reporter construct together with TM. Cells were either incubated with buffer, thrombin (30nM), or a mixture of thrombin (30nM) and Heparin (5 IE; mixture preincubated for 20 minutes) 20 minutes before released AP-activity was measured. Data presented as mean +/- SEM; 3 independent experiments each performed in triplicate.

**Supplementary Fig. S3b**

**Supplementary Fig. S3a**

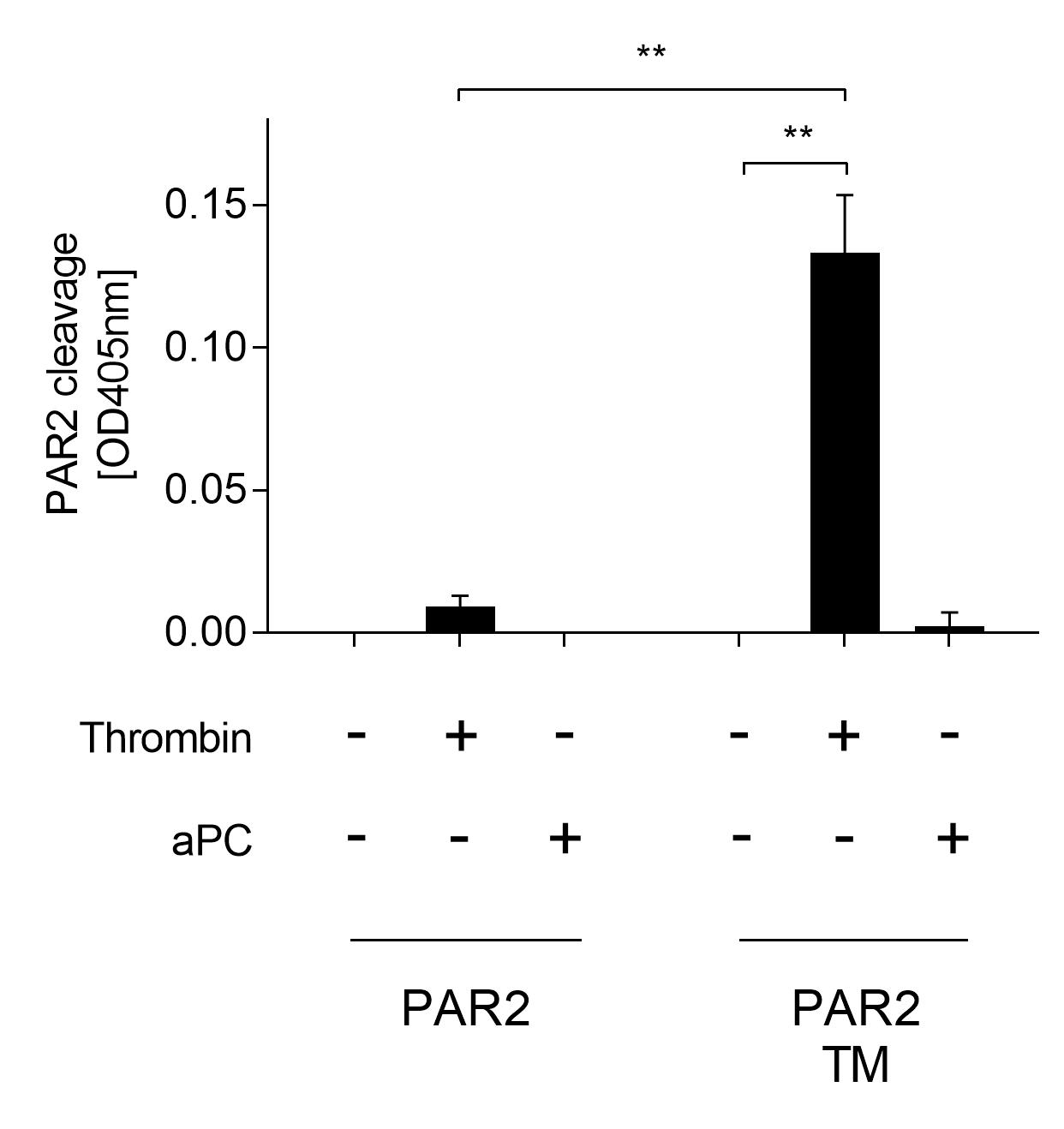

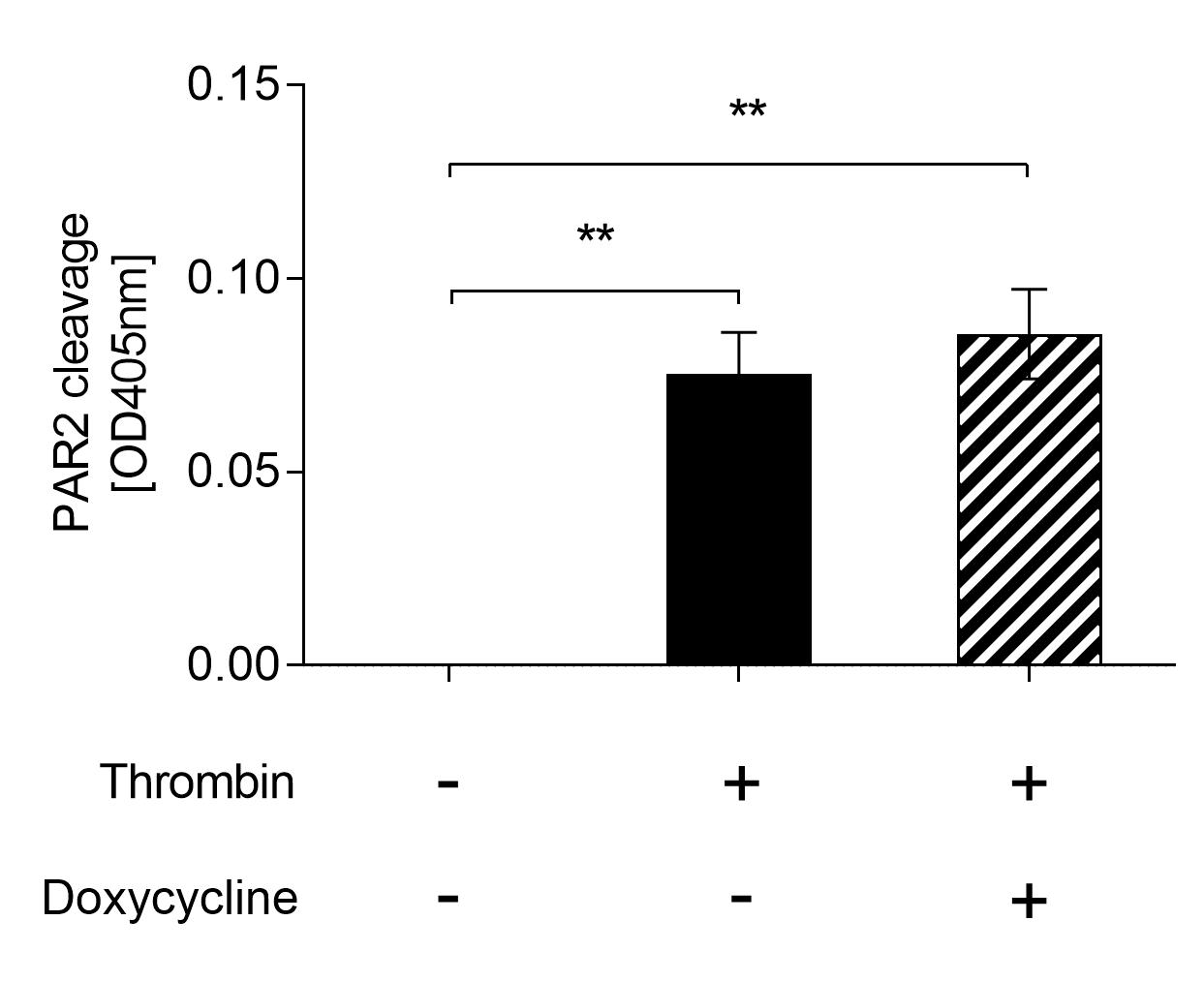

**Supplementary Fig. S3: Testing PAR2 cleavage by other proteases than thrombin.** (**Fig. S3a**) 293T cells transiently overexpressing AP-PAR2 reporter construct either alone or with TM. Cells were incubated with thrombin (30nM) or recombinant aPC (50nM) before PAR2 cleavage was measured. (**Fig. S3b**) AP-PAR2 and TM overexpressing cells were preincubated with the metalloprotease inhibitor doxycycline (o/n; 100µM) before agonist incubation and quantification of PAR2 cleavage. Data presented as mean +/- SEM; 3 independent experiments, each performed in triplicate; ns *p*>0.05, * *p*<0.05, ** *p*<0.01, *** *p*<0.001, using Student *t* test.

**Supplementary Fig. S4a**

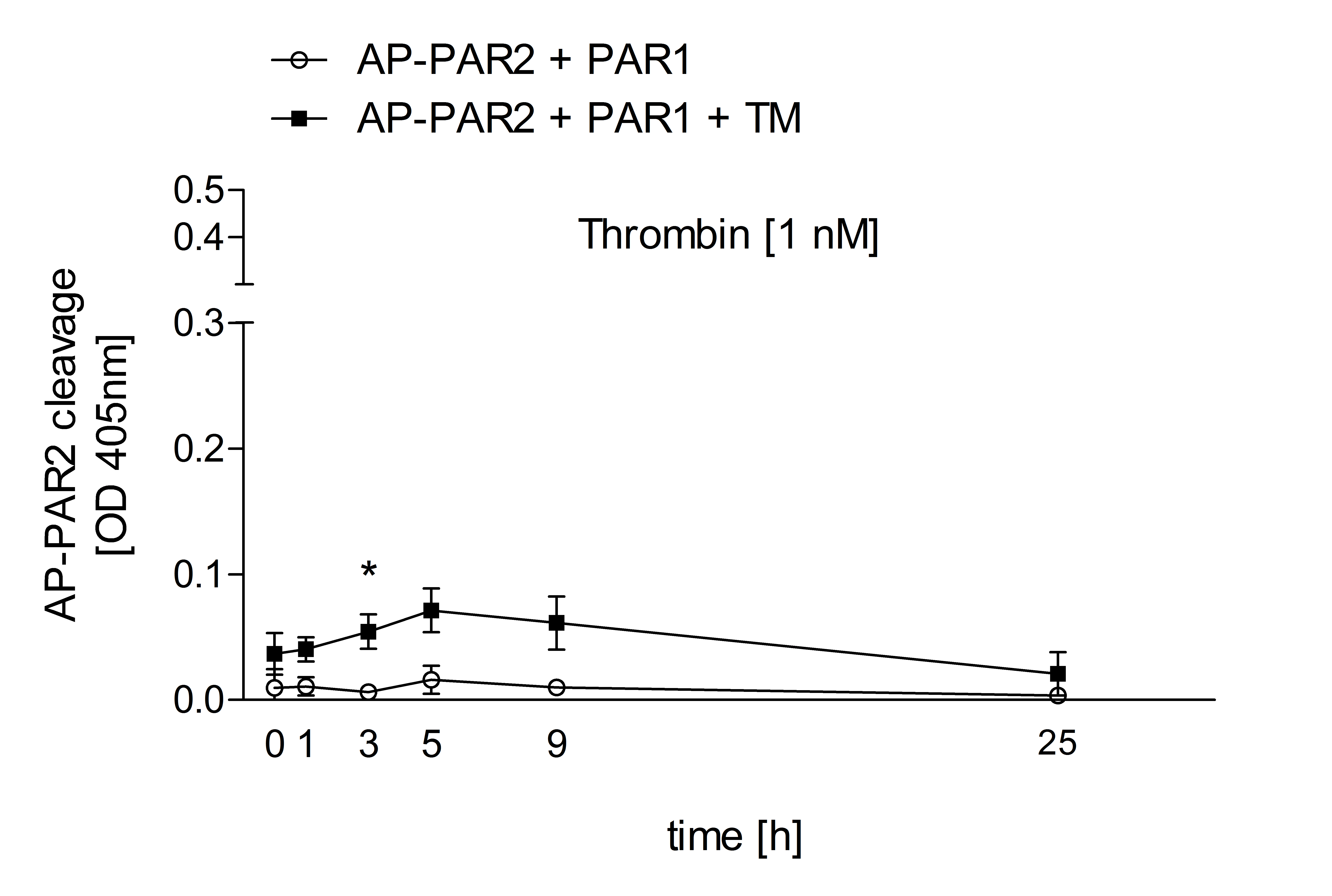

**Supplementary Fig. S4b**

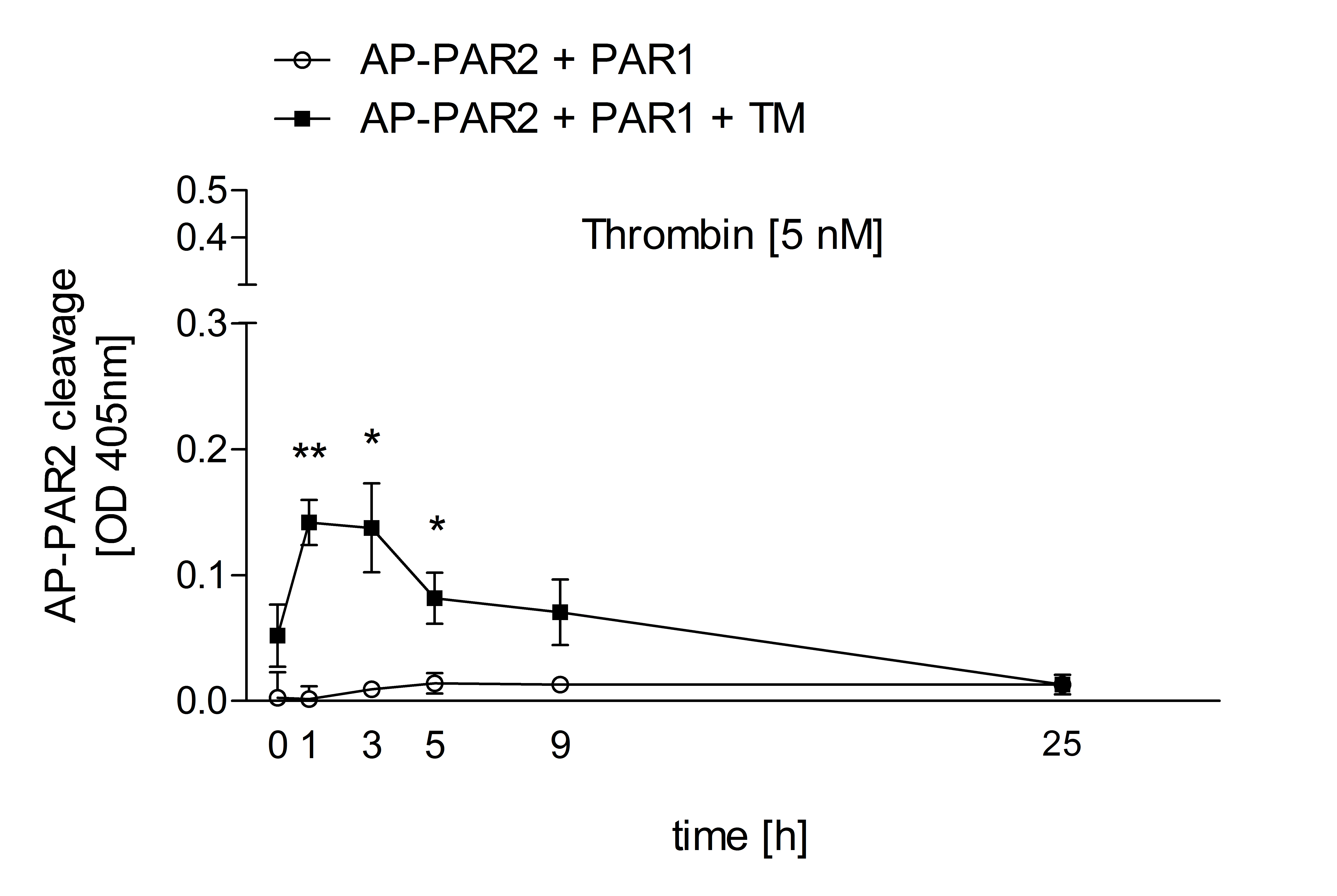

**Supplementary Fig. S4c**

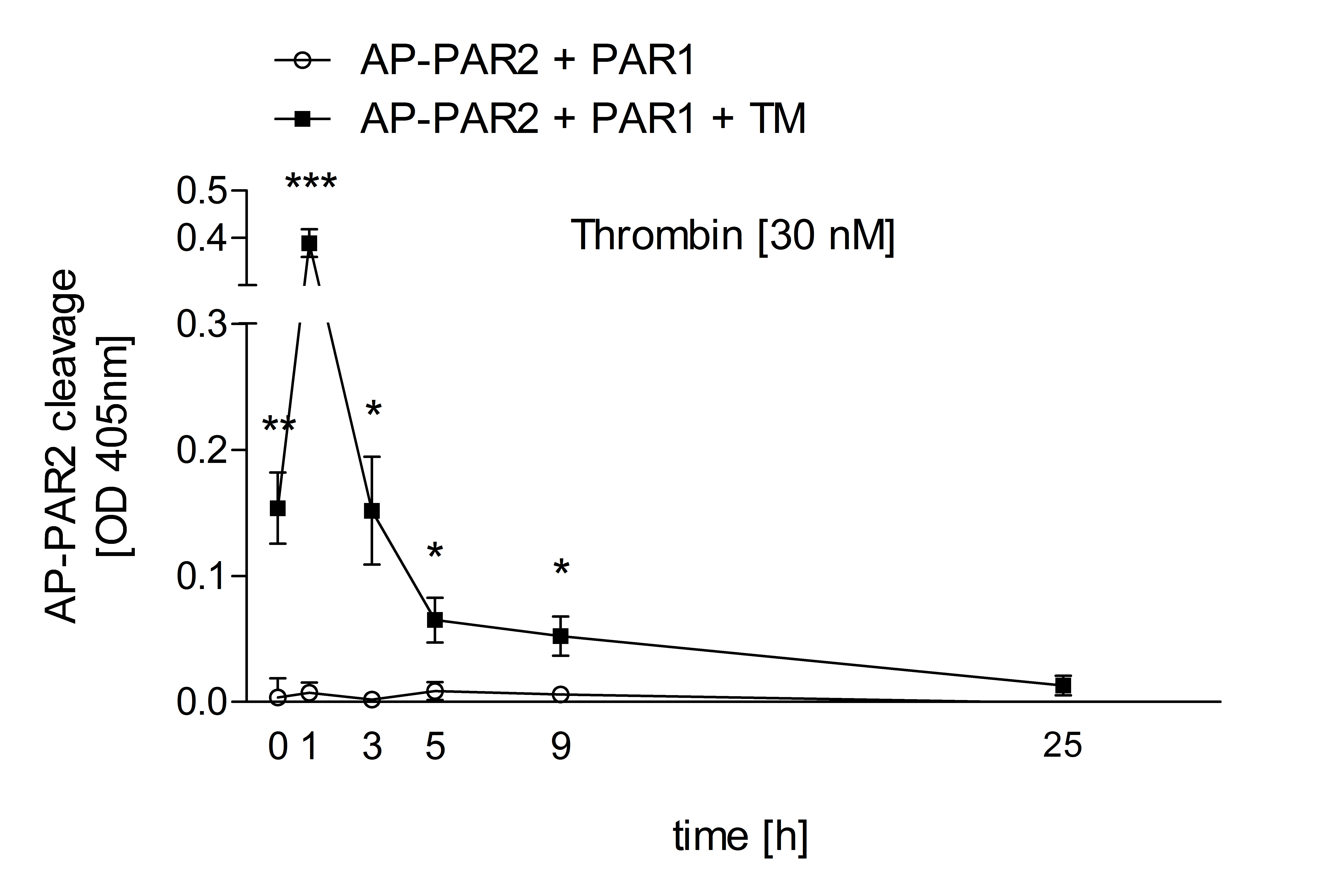

**Supplementary Fig. S4: PAR2 cleavage assay upon prolonged thrombin cleavage.** Cleavage assay of 293T cells overexpressed AP-PAR2 reporter construct together with PAR1 in absence or presence of TM. Cells were incubated over 25h either with buffer or with the indicated concentrations of (**Fig.** **S4a**) thrombin (1 nM), (**Fig.** **S4b**) thrombin (5 nM) and (**Fig. S4c**) thrombin (30 nM). Fresh agonist was added 20 minutes prior to the indicated time point to measure released AP-activity in the supernatant. Stars indicate the comparison of the AP-PAR2 cleavage in cells overexpressing AP-PAR2 + PAR1 to the cleavage in cells overexpressing AP-PAR2 + PAR1 + TM. Data presented as mean +/- SEM; 3 independent experiments each performed in triplicate; ns *p*>0.05, * *p*<0.05, ** *p*<0.01, *** *p*<0.001, using Student *t* test.

**Supplementary Fig. S5**

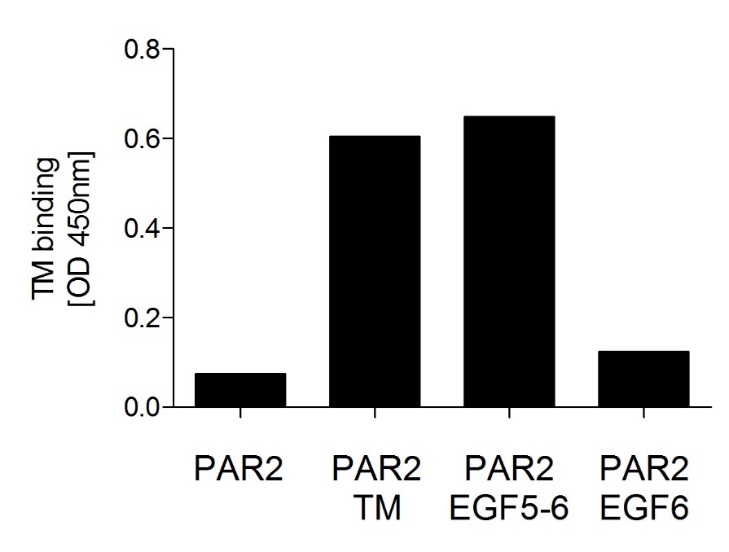

**Supplementary Fig. S5: Quantification of TM expression by cell surface ELISA.** 293T cells transiently expressed AP-PAR2 together with wild type TM (wt), or mutants lacking the lectin domain plus the EGF-like domain 1-4 (TM EGF5-6), or lacking the lectin domain plus EGF-like domains 1-5 (TM EGF6). PFA fixed cells were stained by anti-TM detecting the EGF-like domains 5 and 6 (anti-TM#1009 (141C01(1009)), Thermo Fisher Scientific). Antibody binding was quantified by cell surface ELISA. Data presented as mean +/- SEM; 2 independent experiments in triplicate.

**Supplementary Fig. S6b**

**Supplementary Fig. S6a**

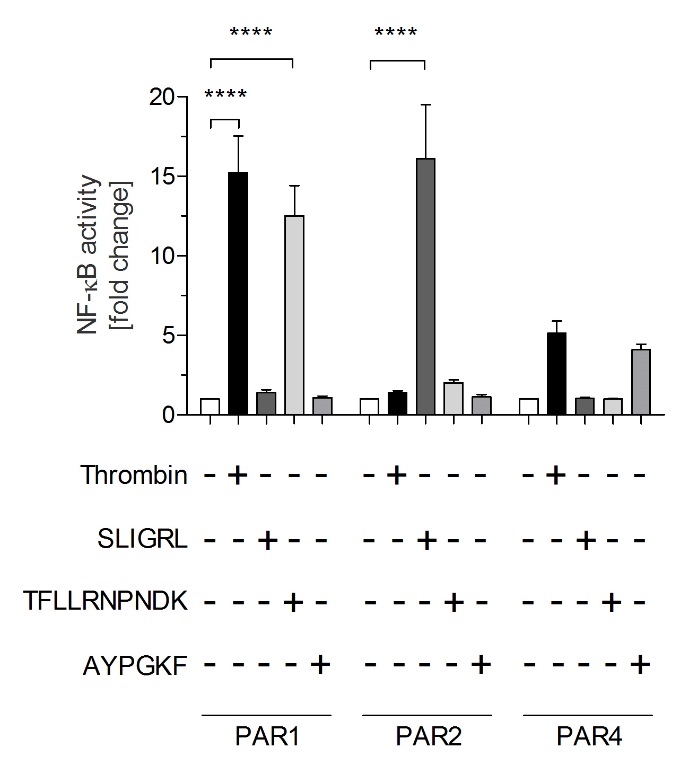

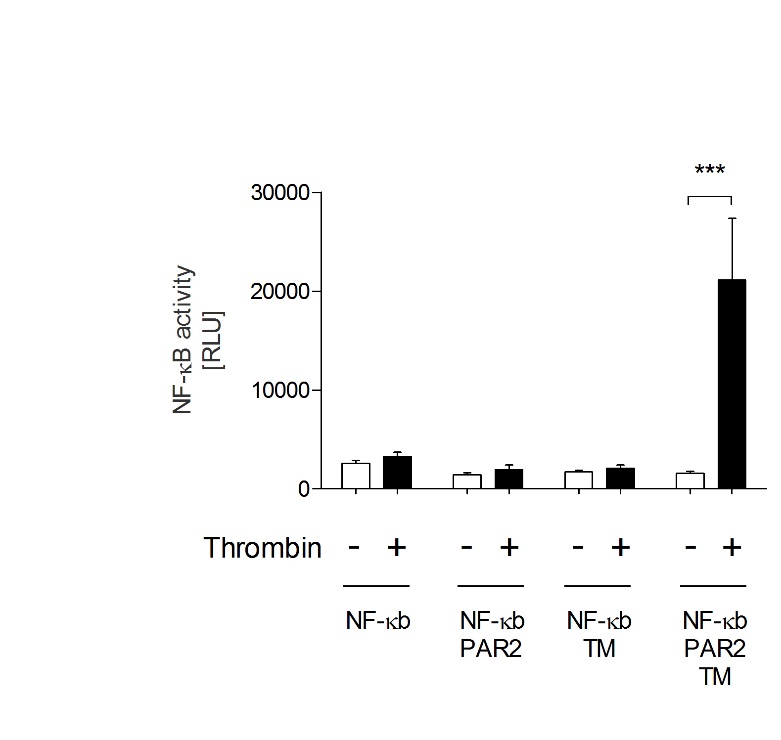

**Supplementary Fig. S6: NF-κB luciferase reporter assay in 293T cells.** (**Fig. S6a**) 293T cells transiently transfected with the NF-κB reporter plasmid pGL4.32[luc2P/NF-κB-RE/Hygro] alone or together with AP-PAR2 and/or TM. Where indicated, cells were incubated with thrombin (30nM). Luciferase activity was quantified using a luminescence plate reader in RLU. (**Fig. S6b**) AP-PAR1, AP-PAR2 or AP-PAR4 constructs were overexpressed together with pGL4.32[luc2P/NF-κB-RE/Hygro] in 293T cells. Transfected cells were induced either with buffer (control), thrombin (30nM), the PAR2 agonist peptide SLIGRL (50µM), the PAR1 agonist peptide TFLLRNPNDK (20µM) or the PAR4 agonist peptide AYPGKF (50µM). Data is shown as fold change of luciferase RLU with control set to 1. Data presented as fold induction of buffer RLU.

Data presented as mean +/- SEM; 3 independent experiments each performed in triplicate; * *p*<0.05, ** *p*<0.01, *** *p*<0.001, **** *p*<0.0001 using 1-way ANOVA.

**Supplementary Table S1**

Sequences of N-terminally truncated variants of TM

| Symbol | | TM variant | Sequence |
| --- | --- | --- | --- |
| 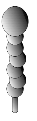 | TM (wild type) | | Consistent with reference sequence NM_00361.2 |
| 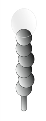 | TM EGF 1-6 | | 5’-GTCGAGCAC*GGATCC*TTCGCGCTC… …TTCCTCTGC*GGATCC*CACTTCCCA-3’ |
| 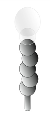 | TM EGF 2-6 | | 5’-GTCGAGCAC*GGATCC*TTCGCGCTC… …TCCTGCACC*GGATCC*GCGACGCAG-3’ |
| 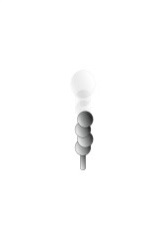 | TM EGF 3-6 | | 5’-GTCGAGCAC*GGATCC*TTCGCGCTC… …CGGTGCGAG*GGATCC*GATGACTGC-3’ |
| 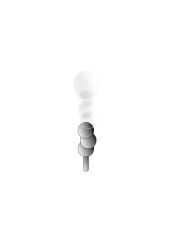 | TM EGF 4-6 | | 5’-GTCGAGCAC*GGATCC*TTCGCGCTC… …GAGTGTGTG*GGATCC*GTGGACCCG-3’ |
| 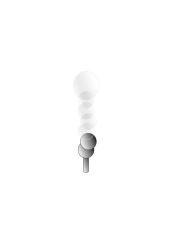 | TM EGF 5-6 | | 5’-GTCGAGCAC*GGATCC*TTCGCGCTC… …CACAGGTGC*GGATCC*TTTTGCAAC-3’ |
| 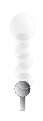 | TM EGF 6 | | 5’-GTCGAGCAC*GGATCC*TTCGCGCTC… …ATCTGCACG*GGATCC*GACGAGTGC-3’ |

TM (wild type) fully matched the entry NM_000361.2 and served as template for mutants. To remove EGF domains, two BamHI sites were introduced (italic) flanking the domains to be removed (boxed). Deletions were obtained by digestion of vectors with BamHI followed by re-ligation allowing to drop domains (box). Scheme of domains provided in Figure 2A, lower panel.

**Supplementary Table S2**

Sequence details of PAR2 construct

**N-terminal linker between AP and PAR2**

pSEAP:	5’-	CCG GGT TAC TCT AGA GTC GGG GCG GCC GGC-3’

Pro Gly Tyr Ser Arg Val Gly Ala Ala Gly_514_

AP-PAR2:	5’-	CCG GGT TAC TCT **G***C***G** **GCC** *CAA GGA ACC AAT* -3’

Pro Gly Tyr Ser **Ala** **Ala** *Gln Gly Thr Asn*

PAR2:	5’-	*TCC TGC AGT GGC ACC ATC CAA GGA ACC AAT*-3’

*Ser Cys Ser Gly Thr Ile Gln Gly Thr Asn*_30_

**C-terminal linker between PAR2 and EGFP**

EGFP:	5’-	ATC GCC ACC ATG-3’

Ile Ala Thr Met

PAR2-EGFP:	5’-	*ACC TCC TAT* ***GGA*** ***TCC*** ATC GCC ACC ATG-3’

*Thr Ser Tyr* **Gly Ser** Ile Ala Thr Met

PAR2:	5’-	*ACC TCC TAT TGA*-3’

*Thr Ser Tyr stop*

The C-terminal sequences of Clontech‘s secretory alkaline phosphatase (pSEAP; GI:2190725) and N-terminal part of Clontech’s enhanced green fluorescent protein (EGFP; Sequence GI:1543070) are underscored. The CDS of PAR2 (homologous to NM_005242.5) is provided (italic), with signal sequence marked (gray) and mutations inserted (bold).

**Supplementary Table S3**

Protein sequence of AP-tagged wt PAR2 reporter construct and mutants

PAR2	S**AA**QGTN	RSSKGRSLIG	KVDGTSHVTG	KGVTVETVFS	VDEFSASVLT	GKLTTVFL

‘PAR2 R36A’	S**AA**QGTN	RSSKGASLIG	KVDGTSHVTG	KGVTVETVFS	VDEFSASVLT	GKLTTVFL

‘PAR2 all-to-A’	S**AA**QGTN	ASSAGASLIG	AVDGTSHVTG	AGVTVETVFS	VDEFSASVLT	GALTTVFL

Position*:	SAA|_27_	|_31_	|_41_	|_51_	|_61_	|_71_

Amino acid sequences of the extracellular N-terminus of PAR2 reporter constructs, with amino acids belonging to the alkaline phosphatase tag (underscored), linker (bold), mutations on PAR2’s N-terminus introduced (box) and first transmembrane domain (grey). *Positions according to UniProtKB‑P55085.

**Supplementary Table S4**

**N-terminal linker between AP and PAR4**

pSEAP:	5’-	CCG GGT TAC TCT AGA GTC GGG GCG GCC GGC-3’

Pro Gly Tyr Ser Arg Val Gly Ala Ala Gly_514_

AP-PAR4:	5’-	CCG GGT TAC TCT AGG GCC *CAG ACC CCC AGC* -3’

Pro Gly Tyr Ser Arg Ala *Gln Thr Pro Ser*

PAR4:	5’-	*AGC CTG TCT GAG AGC GGG CAG ACC CCC AGC* -3’

*Ser Leu Ser Gly Gly Thr Gln Thr Pro Ser* _25_

Sequence details of AP-PAR4 construct

The C-terminal sequences of Clontech‘s secretory alkaline phosphatase (pSEAP; GI:2190725) is underscored. The CDS of PAR4 (homologous to NM_003950.3) is provided (italic), with signal sequence marked (gray).

**Supplementary Scheme S1**

Assembly of PAR2 constructs

| pSEAP | SS | Alkaline Phosphatase | | | Stop |  | |  |  | |
| --- | --- | --- | --- | --- | --- | --- | --- | --- | --- | --- |
| AP-PAR2 | SS | Alkaline Phosphatase | | |  | PAR2 | | Stop |  | |
| PAR2 |  |  | | SS |  | PAR2 | | Stop |  | |
| EGFP |  |  | |  |  | | SS | EGFP | | Stop |
| PAR2-EGFP |  |  | | SS |  | PAR2 | | EGFP | | Stop |

Schematic drawing of the assembly of overexpression constructs composed of signal sequences (SS), the CDS of extracellular alkaline phosphatase, the protease activated receptor 2 (PAR2) with its N-terminal extracellular domain (dashes) and enhanced green fluorescent protein (EGFP) and the stop codon (stop).
